# Dynamic CTCF bridges drive genome organization

**DOI:** 10.64898/2026.05.01.721706

**Authors:** Colleen C. Caldwell, Matilde Sá, Bram J. Hoogland, Andrea Ridolfi, Natal F. Zaghal, Chase P. Broedersz, Andreas S. Biebricher, Gijs J.L. Wuite

## Abstract

Human CCCTC-binding factor (CTCF) is a crucial factor in genome organization, regulating chromatin looping and gene expression. The loop extrusion model (LEM) designates cohesin as the sole active player and limits CTCF to a passive barrier, though emerging evidence suggests a more dynamic role. Using complementary single-molecule techniques, combining dual- and quad-trap optical tweezers, fluorescence microscopy, and atomic force microscopy, we show CTCF forms stable, dynamic DNA bridges. We demonstrate CTCF binds with high affinity and cooperativity, undergoes 1D diffusion, and stiffens DNA. Strikingly, CTCF stabilizes DNA loops without cohesin and forms bridges that slide at low force yet resist rupture at forces exceeding dsDNA stability. These findings suggest a modified LEM in which CTCF directly forms dynamic, force-resistant bridges to stabilize cohesin-established loops, revealing an active role in genome organization.

## INTRODUCTION

The eukaryotic genome is intricately folded within the nucleus, while remaining dynamically accessible for gene expression, and DNA replication, recombination, and repair.^1–6^ The LEM has emerged as the central framework for understanding how interphase genome organization is established.^7^ In this model, the loop extruder cohesin initiates and expands chromatin loops until encountering convergent CTCF located at CTCF binding sites (CBSs) (Figure 1A), providing an explanation for the “CTCF convergence rule” and loop patterns derived from chromatin capture data and simulations.^8–10^ Single-molecule studies of cohesin have contributed significantly in advancing our understanding of mechanisms underlying chromatin loop formation.^11,12^ In contrast, the exact role of the other central player in chromatin organization, CTCF, remains mechanistically obscure despite being an essential protein across bilaterians.^13^ Beyond chromatin loop formation, CTCF performs roles in alternative splicing, X chromosome inactivation (XCI), and DNA repair, making its context-dependent mechanisms of action of particular interest.^14–18^

**Figure 1.**
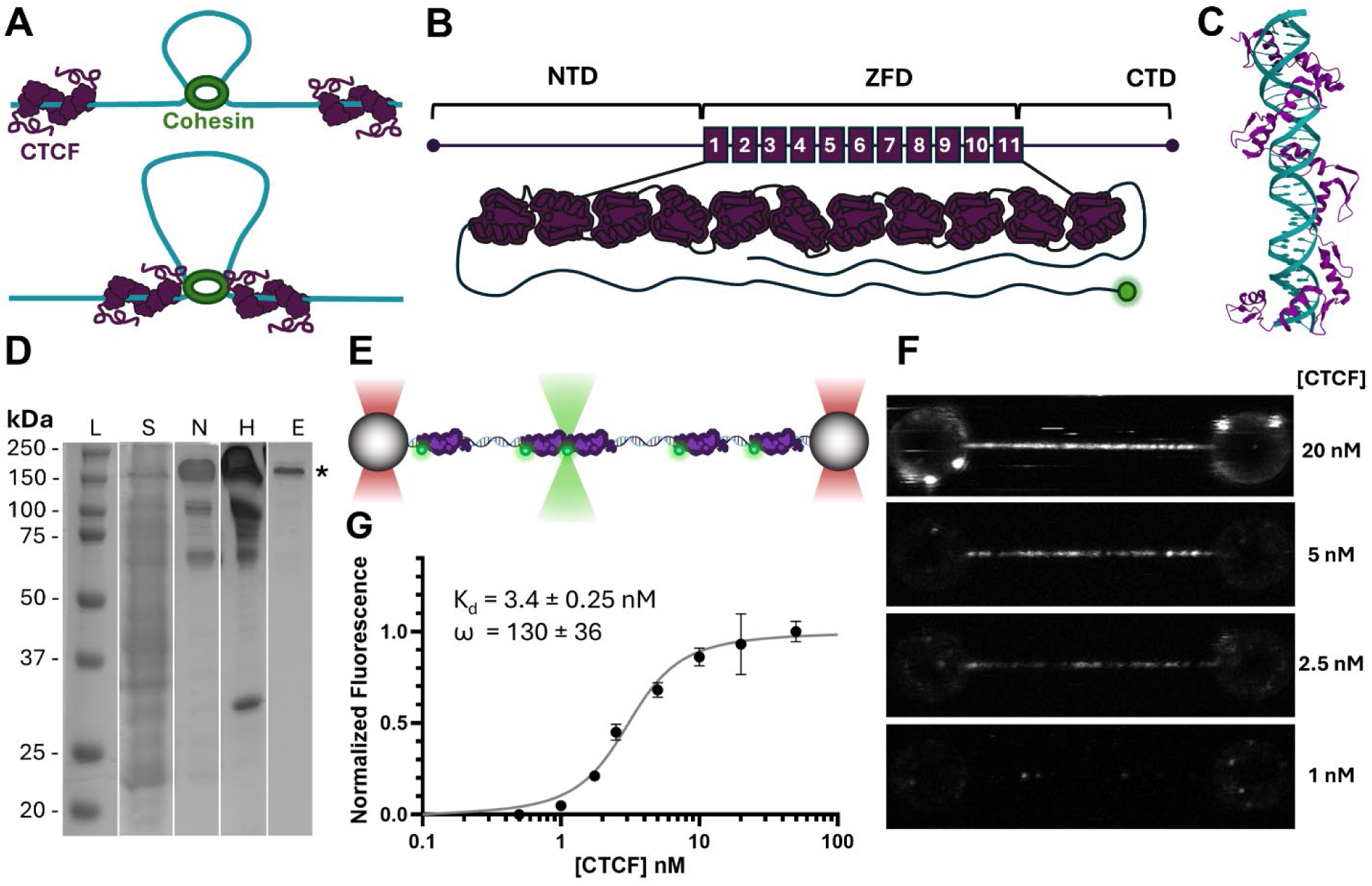
Full-length CTCF binds non-specific DNA with high affinity and positive cooperativity. (**A**) Illustration of the loop extrusion model highlighting the role of cohesin in expanding loops and CTCF as a barrier. (**B**) Schematic of full-length CTCF domains: N-terminal domain (NTD), a central zinc finger domain (ZFD) comprising 11 tandem C2H2 zinc fingers, and a C-terminal domain (CTD). N-terminal labelling with Alexa Fluor 555 was employed. (**C**) Crystal structure of CTCF ZF3-11 binding the major groove following the DNA helix (PDB:8SSQ)^20^. (**D**) SDS-PAGE analysis verifying purification of CTCF. Lanes include (L) molecular weight markers, (S) supernatant, (N) nickel column elution, (H) heparin column elution, and (E) size exclusion elution samples. See also Figure S1. (**E**) Illustration depicting CTCF^555^ binding to tethered λ-DNA and imaging using confocal scanning. (**F**) Selected representative confocal fluorescence images of DNA incubated with increasing concentrations of CTCF^555^ (500 pM to 50 nM). At low concentrations, sparse binding is observed, while higher concentrations result in saturation of DNA-bound CTCF. (**G**) Quantification of total fluorescence associated with DNA across replicates (N ≥ 5 DNA molecules) as a function of CTCF concentration. Data are fit to the McGhee-von Hippel model, revealing an effective dissociation constant (K_d_) of 3.4 ± 0.25 nM and a cooperativity factor (ω) of 130 ± 36, indicating positive nearest-neighbor cooperativity in CTCF binding to DNA. Errors are S.E.M.

The multifunctionality of CTCF may stem, in part, from its unique structural composition (Figure 1B). The central region of CTCF contains 11 tandem C2H2 zinc fingers (ZFs),^19,20^ the α-helix of each lying within the major groove of DNA, driving CTCF to form a coil around the DNA helix (Figure 1C).^20^ Flexible linkers connect the tandem ZFs and allow CTCF to adopt variable conformational states.^21^ Early investigations suggest this flexibility may permit alternative ZF engagement and allow CTCF to act as a multivalent transcription factor.^19^ More than half of CTCF consists of the disordered N- and C-termini.^22^ Both termini have been identified as playing a role in CTCF oligomerization^13,23–25^ along with the ZF domain^20,26^ while the N-terminus mediates interaction with cohesin.^27^ Recent single-molecule work suggests that this interaction with cohesin modulates or halts its loop extrusion activity.^28^

Since the discovery of CTCF in 1990 as a transcription factor,^29^ several major shifts have occurred in its identified role, from enhancer regulation^30^ to global genome organizer.^14^ While recent understanding of CTCF focuses on halting of cohesin loop extrusion, earlier models proposed that CTCF may function in part by directly bridging DNA itself.^31^ Particularly strong evidence for a role in bridging beyond its interplay with cohesin lies in the discovery of trans-chromosomal interactions between CTCF bound on different chromosomes, which cannot be explained by loop extrusion.^32,33^ Additionally, the function of CTCF appears to extend beyond binding at specific sites, as depleting CTCF results in greater perturbations than removing CBSs.^34^ This supports the relevance of CTCF binding on non-specific DNA beyond previously suggested facilitated diffusion.^12,35,36^

Here we combine dual- and quad-trap optical tweezers with fluorescence microscopy to characterize CTCF’s interaction with DNA, revealing direct stabilization of DNA loops via bridging. We find that these dynamic bridges slide along non-specific DNA at low force yet resist rupture beyond the stability of dsDNA. Monitoring sliding bridges provides a quantitative measurement of the friction force of a protein sliding along DNA. Revealing the role of CTCF on non-specific DNA and resolving the long-standing question of CTCF’s ability to independently bridge DNA is critical for developing models that account for all active components.

## RESULTS

### CTCF binds cooperatively and slides on non-specific DNA

To investigate the behavior of CTCF in a form that reflects its native structure and function, we expressed and purified the full-length protein with an N-terminal 6X-histidine tag in *Escherichia coli*, preserving all 11 ZFs and the disordered termini, which make up over 60% of the protein (Figures 1B and 1D). The protein was then purified utilizing a combination of affinity and size-exclusion chromatography (Figure S1). For fluorescence studies, the N-terminal amine of the purified CTCF was labelled directly with Alexa Fluor 555-NHS (CTCF^555^), enabling single-molecule visualization without addition of bulky, disruptive tags.

Using dual-trap optical tweezers combined with fluorescence microscopy, we investigated the binding of CTCF^555^ to non-specific dsDNA (λ-DNA, ∼16.5 µm) tethered between optically trapped beads (Figure 1E). DNA molecules were maintained under constant tension to ensure extension and prevent spontaneous loop formation. dsDNA molecules were incubated with increasing concentrations of CTCF^555^ (0.5 to 50 nM) and fluorescence signal along the dsDNA was recorded (Figure 1F). As CTCF concentration increased, binding became progressively denser until a maximum intensity was reached, consistent with saturation of available binding sites. Using the measured brightness of a single monomer (Figure S2), we calculated the total number of bound monomers and estimated a binding footprint of 29 ± 1.6 bp. Total fluorescence signal was normalized and plotted as a function of CTCF concentration (Figure 1G), and the resulting binding curve fit to the McGhee-von Hippel model.^37^ This analysis revealed a dissociation constant (K_d_) of 3.4 ± 0.25 nM and nearest neighbor cooperativity factor (ω) of 130 ± 36, indicating a remarkably high affinity for non-specific dsDNA and positive binding cooperativity. While our observed affinity for non-specific dsDNA is much higher than reported in previous studies,^21,35,38^ we propose that this could be explained by our use of a non-truncated protein combined with a much longer non-specific dsDNA substrate. On the other hand, our finding of a positive cooperativity aligns with evidence from cellular contexts. The efficiency of CTCF-dependent insulation is known to be enhanced by tandem CBSs, suggesting that clustered CTCF molecules act cooperatively to establish and maintain TAD boundaries^39–41^.

Incubating dsDNA at sub-nanomolar CTCF^555^ concentrations enabled us to follow individual dsDNA binding events in real time (Figure 2A) and to differentiate between mono- and oligomeric binding of CTCF (Figure S2). In addition to binding and dissociation, the resulting kymographs reveal that CTCF undergoes 1D diffusion (“sliding”) along the dsDNA molecule (Figure 2B). Monomeric CTCF^555^ traces (selected based on intensity, Figure 2B) were tracked and the resulting trajectories analyzed to calculate mean squared displacement (MSD; Figures 2C-D). A linear fit of MSD data pooled over all traces versus time lag yielded a diffusion constant (D) of 0.092 ± 0.0025 μm^2^/s (Figure 2D), which is comparable with rotationally-coupled diffusion by proteins similar in size to CTCF (Supplemental Note 1). This is not unexpected for a protein featuring specific binding^42^ and is in agreement with previous *in vitro* studies.^12^

**Figure 2.**
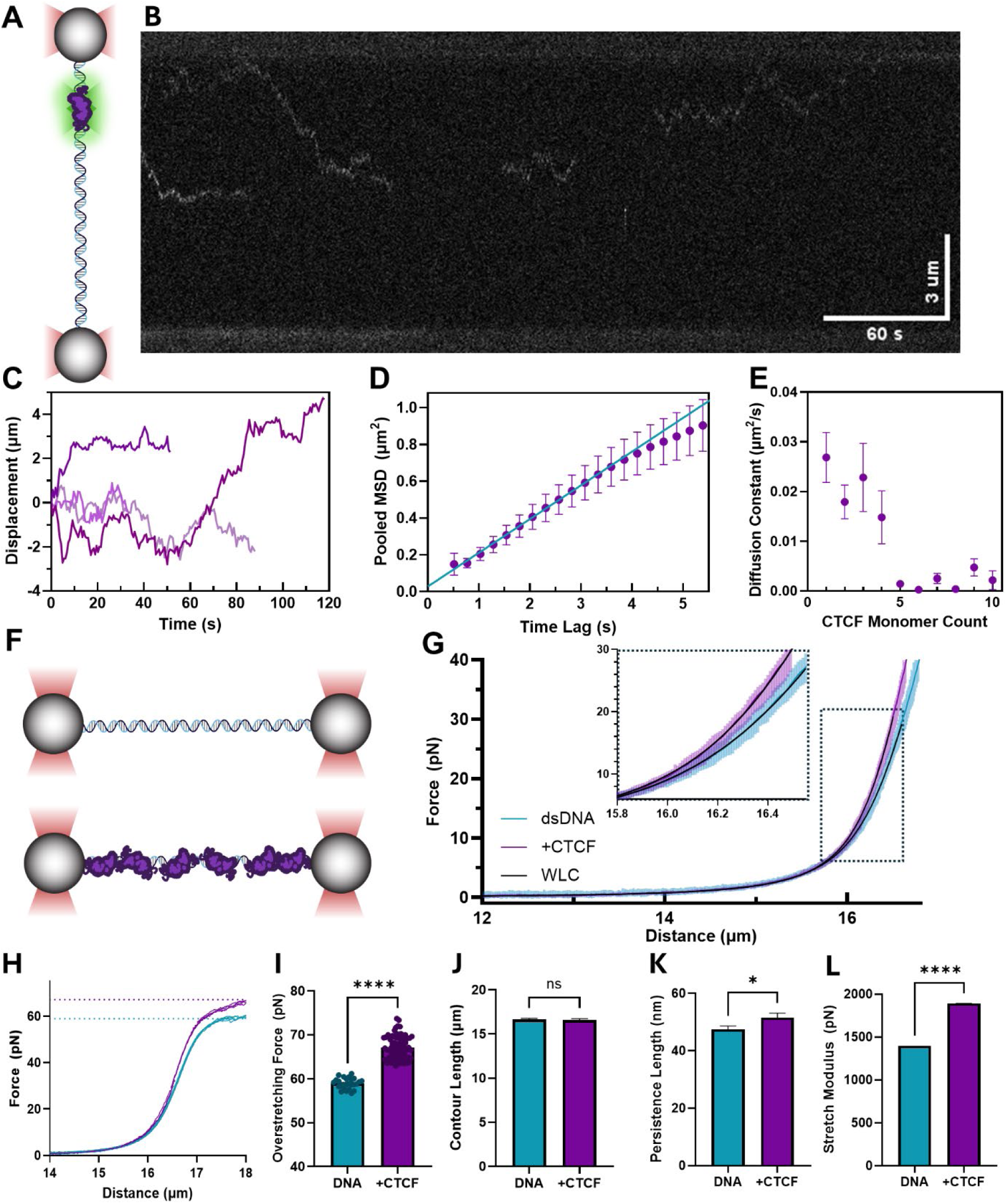
CTCF undergoes 1D diffusion on non-specific DNA and alters the mechanical properties of dsDNA. (**A**) Schematic illustrating tracking of CTCF binding and diffusion along tethered DNA using dual-trap optical tweezers paired with confocal microscopy. (**B**) Representative kymograph showing the binding, sliding, and dissociation of mobile individual CTCF molecules along DNA over time. The horizontal axis represents time, and the vertical axis represents position along the DNA. (**C**) Individual representative trajectories of diffusing CTCF particles extracted from kymographs. (**D**) Mean squared displacement (MSD) as a function of time lag for pooled unconstrained, monomeric CTCF trajectories (N = 17 monomers). The diffusion constant, 0.092 µm^2^/s, is calculated from the linear fit. (**E**) Average diffusion constants of all CTCF trajectories sorted by oligomeric state. (**F**) Schematic of dual-trap optical tweezers setup used to measure force-extension curves of bare DNA (top) and DNA saturated with CTCF (bottom). (**G**) Force-extension curves for bare DNA (cyan) (N = 57 molecules) and CTCF-coated DNA (purple) (N = 30 molecules). The dashed black line indicates a fit to the extensible worm-like chain (WLC) model. The inset highlights the deviation between bare and CTCF-coated DNA at higher forces. Error bars show the range in measured values. (**H**) Overstretching behavior of bare DNA (N = 25 molecules) and CTCF-coated DNA (N = 30 molecules). The force at which the B-to-S DNA transition occurs is elevated in the presence of CTCF. (**I**) Overstretching force significantly increases for CTCF-coated DNA (****P = 0.0001). (**J**) Contour length is unchanged for CTCF-coated DNA (ns). (**K**) Persistence length increases upon CTCF binding, indicating reduced flexibility (*P = 0.05). (**L**) Stretch modulus is significantly higher for CTCF-coated DNA, suggesting greater stiffness (****P < 0.0001). Error bars represent S.E.M. Statistical significance was determined using unpaired t-tests.

In addition to the population of mobile CTCF^555^ monomers, more complex diffusion behavior was observed, including immobile monomers and transitions in mobility such as pausing, while oligomer binding events showed a tendency toward slower diffusion (Figure 2E). The latter observation is of interest since it would be in agreement with a scenario where the ZF-domain interacts with the DNA while N- and C-termini would be accessible to establish protein-protein interaction in a way which allows multiple proteins to simultaneously interact with the same DNA.

### CTCF binding alters DNA response to force

To assess how CTCF binding alters the mechanical properties of dsDNA, we measured the force-extension behavior of single dsDNA molecules before and after coating with CTCF under saturating conditions and compared the resulting force-extension curves (Figure 2F-G). The comparison revealed several CTCF-induced alterations of dsDNA mechanics, notably an increased onset force of the overstretching plateau from 59 ±1 pN for bare dsDNA to 67± 3 pN for dsDNA coated with CTCF, which suggests stabilization of duplex DNA against force-induced melting (Figure 2H). A more detailed quantification is enabled by fitting force-extension curves of dsDNA with the extensible worm-like-chain (eWLC) model in the absence and presence of CTCF (Figure 2G). Changes to the three fit parameters revealed the following tendencies. The contour length (L_C_) of the dsDNA was not significantly altered, suggesting that CTCF does not shorten or lengthen DNA upon binding (Figure 2J). A modest increase in the persistence length (L_P_) (from 47 ± 1 to 52 ± 2 nm) occurred following CTCF binding, indicating that CTCF reduces dsDNA bending flexibility (Figure 2K). The most substantial change was in the stretch modulus, which increased significantly by 36% (from 1400 ± 100 to 1900 ± 100 pN) for CTCF-coated dsDNA, consistent with a higher resistance of dsDNA against stretching under force (Figure 2L). These effects are indicative of the formation of a more rigid dsDNA structure in the context of the CTCF:dsDNA complex.

The mechanical effects of CTCF on dsDNA are consistent with the reported structures of CTCF bound to its target sequence, in which the ZFs coil around the dsDNA helix.^20^ Such a structure would explain the increased DNA resistance against deformation as well as its reduced bending when bound to CTCF. While many proteins are known to alter dsDNA structure and mechanics, it is somewhat unexpected that the same protein displays 1D diffusion along the DNA. However, this combination of features is not unprecedented, as similar characteristics are known for the mitochondrial transcription factor (TFAM), which not only shows diffusion while affecting the DNA mechanics but also displays cooperative binding with oligomer formation.^43^

### CTCF stabilizes DNA loops in the absence of a loop extruder

To test whether CTCF promotes or stabilizes loops independently of cohesin, we employed AFM and dual-trap optical tweezers as complementary assays (Figure 3A-B), which allowed us to directly visualize and mechanically probe the impact of CTCF on DNA topology. AFM imaging was used to directly visualize dsDNA loops stabilized by CTCF. This was accomplished by quantifying the number of intra-strand crossings of a linear 2.1 kbp plasmid in the absence and presence of 36 nM CTCF. These results demonstrated that CTCF-bound dsDNA displayed a much higher fraction of intra-strand crossings (85 ± 3%) compared to bare DNA (12 ± 3%, Figure 3C-E). The end-to-end distance also decreased in dsDNA molecules following addition of CTCF, consistent with CTCF-induced stabilization of looped structures (Figure 3F).

**Figure 3.**
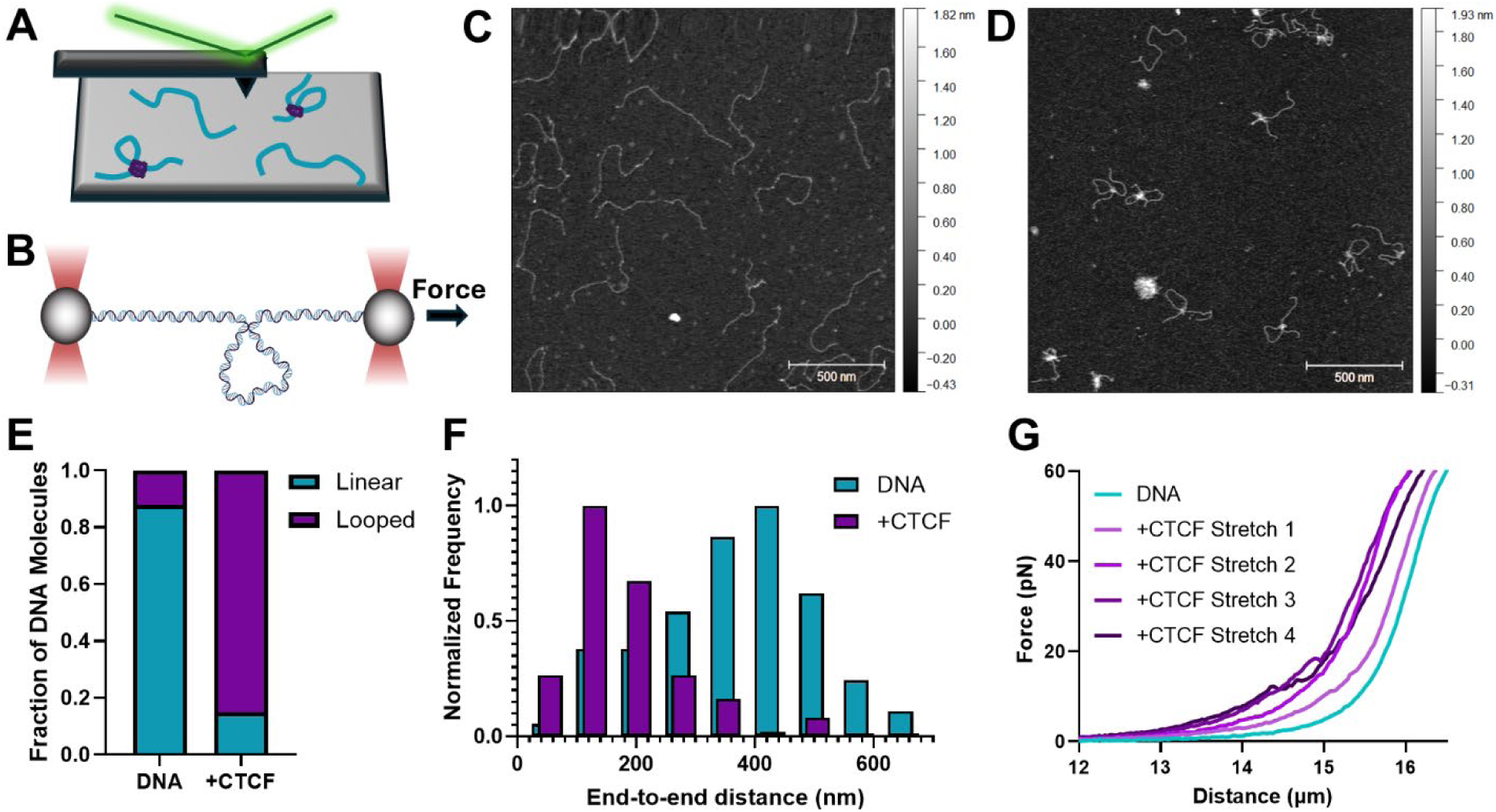
CTCF promotes loop formation in the absence of a loop extruding partner. (**A**) Schematic of the AFM setup used to image DNA and CTCF-mediated loops. (**B**) Schematic of dual-trap optical tweezers setup used to probe CTCF-mediated DNA loops allowed to form on relaxed DNA (**C**) Representative AFM image of bare DNA molecules, which predominantly adopt a linear conformation. (**D**) Representative AFM image of DNA molecules incubated with CTCF, showing an increase in looped DNA structures. (**E**) Quantification of DNA topology from AFM images. Bare DNA predominantly adopts a linear conformation (88 ± 2.7%, N = 144 molecules), while CTCF-bound DNA shows a significant increase in looped structures (85 ± 3.2%, N = 126 molecules). Indicated errors denote standard error of the proportion. (**F**) Normalized distribution of end-to-end distances for bare DNA and CTCF-bound DNA. CTCF binding reduces the end-to-end distance, consistent with DNA compaction and loop formation. (**G**) Force-distance response following formation of CTCF-mediated loops on relaxed DNA resulting in heterogenous shortening.

To further investigate the mechanical properties of the CTCF-dependent loops observed by AFM, we employed the previously described dual-trap optical tweezers assay. In contrast to the earlier experiments performed under tension, we significantly reduced bead separation during CTCF-incubation (10 µm), providing sufficient slack to enable stochastic loop formation (Figure 3B). Subsequent force-extension curves revealed variable shortening of the DNA bound by CTCF compared to bare controls, consistent with the stabilization of looped configurations (Figure 3G). While application of higher forces to shortened constructs resulted in some lengthening, no significant rupturing events were observed even at the highest attainable forces (> 60 pN), and the DNA remained shorter. These findings indicate that the loops are stable yet potentially dynamic, slipping under tension.

### CTCF bridges slide in response to force and resist rupture

While AFM and dual trapping gave clear indications of CTCF-induced loop stabilization, these techniques did not allow for control of either loop formation or loop probing. We therefore switched to a quad-trapping OT set-up, which allows for highly controlled manipulation of four beads simultaneously.^44–46^ This not only permitted us to form CTCF-induced DNA bridges in a highly controlled manner, but to subsequently probe the formed protein bridge using two distinct configurations of the bridged DNA strands (Methods and Figures S4-5). These approaches enabled us to separately interrogate the stability of the protein-protein interaction forming the bridge (Figure 4A) or the protein-DNA interaction of the bridging proteins with their respective DNA strand (Figure 4B).

**Figure 4.**
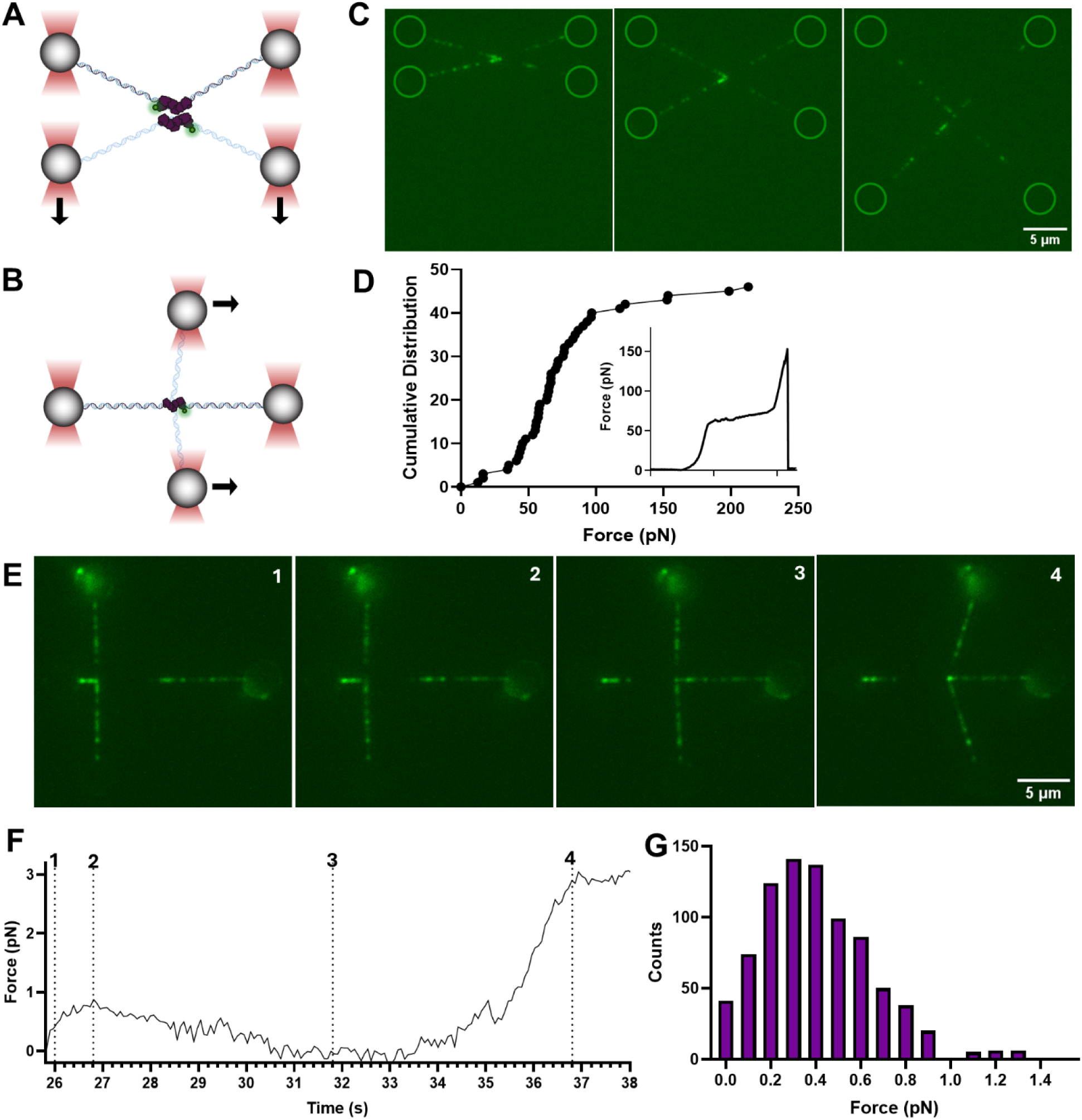
CTCF-mediated bridges slide readily with applied force and resist rupturing at high forces. (**A**) Schematic of quad-trap optical tweezers setup used to measure rupture forces of CTCF-mediated DNA bridges. DNA molecules tethered between optically trapped beads were crossed and incubated with CTCF to form a bridge. Force was applied by increasing distance between bridged DNA molecules. (**B**) Schematic of quad-trap optical tweezers setup used to measure forces under which CTCF-mediated DNA bridges slide. Force was applied by sliding the vertical strand along the horizontal strand. (**C**) Representative widefield fluorescence microscopy images of a CTCF-mediated DNA bridge. Green circles highlight bead locations. (**D**) Cumulative distribution of rupture forces for CTCF-mediated bridges (N = 46 ruptures). The inset shows a representative force-extension curve with a sharp rupture event at high force. Most bridges resist rupture forces exceeding those required for DNA overstretching (> 65 pN). (**E**) Representative widefield fluorescence microscopy images of a CTCF-mediated DNA bridge sliding in response to force application. (**F**) Force exerted on the CTCF mediated bridge over time as the bridge slides along DNA. Marked time points (dotted lines) correspond to the preceding images. (**G**) Histogram of forces measured during mobile sliding of CTCF bridges (N = 16 bridges). The majority of sliding occurs below 1 pN.

To generate CTCF-mediated DNA bridges, two λ-DNA molecules were tethered between separate pairs of optically trapped beads and positioned at a right angle to each other to create a cross-like configuration, such that the two dsDNA strands make contact at their intersection (Figure 4B). This geometry promoted the formation of DNA bridges upon moving into a channel containing 25 nM CTCF⁵⁵⁵. Following incubation, the construct was transferred back into a buffer-only channel for DNA manipulation and imaging (Figures S4-5).

Fluorescence intensity analysis allowed us to determine that the formed bridges did not feature single CTCF molecules, but were composed of oligomers, mainly dimers and trimers (Figure S3). These results suggest that CTCF-mediated bridges require a CTCF molecule bound to each of the dsDNA strands, which subsequently form a protein-protein connection with each other. CTCF-mediated bridges also form with notable efficiency. 83% of attempts to form bridges in the presence of CTCF (*N* = 40) resulted in stable bridging between DNA strands.

Next, we set out to probe the stability of the protein-protein interactions mediating the bridge using the instrument’s capability to move two traps simultaneously. To this end, we oriented the two bridged strands roughly parallel to each other and then moved one strand further away from the other one (Figure 4A and Supplementary Video 2). This configuration ensures that tension is applied at a right angle to the DNA, which enables probing of the protein-protein bridge stability isolated from other factors. Notably, the majority of bridges displayed a remarkable stability and thus remained intact even as the applied force exceeded 65 pN—the threshold for overstretching B-form DNA (Figure 4C). In most cases, the DNA tether failed before the bridge itself (Supplementary Video 3), therefore, the resulting cumulative force distribution (Figure 4D) reflects only a lower bound of the stability of the protein-protein bridge which may greatly exceed 65 pN. To probe the resistance of CTCF-mediated bridges to dragging forces, we employed an orthogonal DNA configuration where one strand was dragged along the axis of the other, causing the bridge to slide (Figure 4B). In striking contrast to the high force resistance of the protein-protein bridges to rupturing, CTCF complexes involved in the bridge retained a remarkable motility on the dsDNA and were readily dragged along the DNA axis when the other strand was moved. Fluorescence imaging showed that the bridges remained tethered and moved in the direction of the applied force, often displacing other CTCF molecules along the DNA (Figure 4E and Supplementary Video 1). These features were observed independently of which strand was moved and the direction of movement, showing that bridge mobility is not dependent on directionality. The force applied to the bridge was determined by measuring the force on the stationary bead as the bridge was moved and subtracting the tension on the tethered dsDNA (Figure 4E). In most cases (13/16), an initial peak in force was observed at the onset of sliding, followed by a phase of free sliding, during which the average force dropped to 0.4 ± 0.1 pN and rarely exceeded 1 pN (Figure 4F-G), until sliding bridges eventually encountered a DNA obstacle at which point movement ceased. Notably, the observed dragging forces can be related to and are in remarkable agreement with the observed CTCF diffusion coefficient deduced above (Supplemental Note 1 and 2).

CTCF-mediated bridging behavior is likely driven by protein-protein interactions, as bridges were always observed to be oligomers and never monomers, suggesting at least two proteins are required for bridging. The stability of the CTCF-mediated bridges against rupture forces is remarkably high, exceeding the stability of the dsDNA (∼60-70 pN) it is bridging. The resistance against rupturing of bridges stabilized by CTCF is also notably higher than other observed DNA bridging proteins.^47–49^ This is even more remarkable when contrasted with the much lower forces (< 1 pN) at which displacement of the CTCF bridges is induced by dragging it over the DNA. While remarkable, these features are not entirely unprecedented, as strikingly similar XLF:XRCC4 “sliding sleeves” also show high resistance to rupture while easily sliding along dsDNA.^44^ Similar to the sliding sleeves, CTCF coils around dsDNA, potentially explaining their high resistance to being pulled off of DNA. In addition to resisting displacement from DNA, the protein-protein interaction must also have notable resistance. The N- and C-termini, along with CTCF ZFs, have separately been found to play roles in oligomerization,^20,26,50,51^ perhaps additively explaining the notable stability of CTCF bridges.

## DISCUSSION

The development of the LEM relied on essential single-molecule observations of loop extruders in action,^52,53^ but similar characterization of CTCF has remained a persistent barrier. We present an in-depth single-molecule study of CTCF and demonstrate that CTCF binds to long, non-specific dsDNA with high affinity, exhibits positive cooperativity, forms oligomers, and stiffens bound DNA. Notably, we find CTCF stabilizes loops by bridging dsDNA in the absence of cohesin. Furthermore, since CTCF has been implicated in processes beyond chromatin organization – from sister chromatid cohesion^54,55^ to DNA repair^16,17^ – gaining mechanistic understanding becomes still more critical in determining its role in these biological processes.

Our results give structural insights into the CTCF:DNA complex, specifically in the context of non-consensus binding, where no structures currently exist. Notably, the coiled structure of CTCF (Figure 5A) reported on consensus DNA^20^ fits with the behaviors we observe on non-specific DNA. First, the rotationally coupled diffusion of CTCF is consistent with the coiled ZFs of CTCF tracking the major groove (Figure 5B). Furthermore, we observe mechanical impacts of CTCF association with DNA consistent with the coiled structure: inhibition of force-induced melting, increased resistance to extension, and reduced flexibility. Together, this suggests the formation of a CTCF-dsDNA complex in which CTCF forms a coating around dsDNA that increases the DNA resistance against deformation (Figure 5C). While the ZFs are implicated in DNA binding, this leaves the disordered termini free to engage in protein-protein interactions. Accordingly, we observe CTCF oligomerization in line with prior work suggesting the termini as drivers of these interactions.^13,23,24^ In the context of the CTCF-mediated bridging and looping that we observe, interacting molecules are not constrained to termini-to-termini interactions. Direct contacts between ZFs are also plausible in bridging orientations (Figure 5D). ZFs have been previously implicated in oligomerization^20,26^ and such contacts could contribute to the bridges’ high force resistance while still accommodating sliding. This rationale is analogous to interactions between parallel sleeves of XLF-XRCC4, which form sliding bridges that resist rupture.^44^

**Figure 5.**
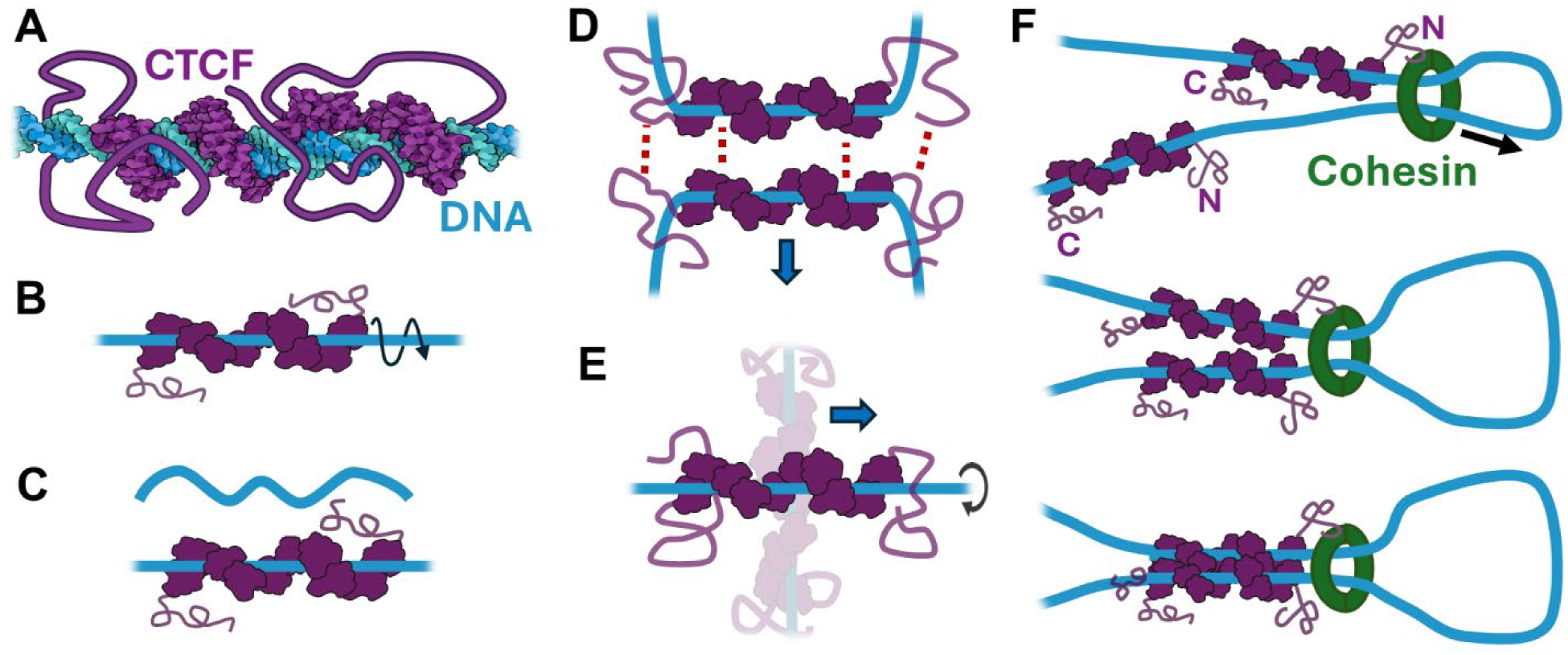
Schematic representation of observed CTCF features. CTCF is illustrated in purple and DNA in blue. (**A**) Illustrative model of CTCF (purple) bound to the DNA helix (blue) with ZFs sitting within the major groove, resulting in a coiled protein structure that fully wraps the DNA. Illustrative model generated using AlphaFold. Our observation of rotationally coupled diffusion (**B**) and reduction in DNA malleability upon CTCF binding (**C**) align with the structure of CTCF coiled around the DNA helix. (**D**) The ability of CTCF-bridges to resist rupturing at forces beyond the stability of DNA may be understood due to the structure of CTCF encircling the DNA helix along with several points of protein-protein interaction. (**E**) During dragging of CTCF-bridges the protein is constrained and unable to rotate. Instead, the DNA, which has free ends, may rotate in response to translocation of the bridge (see supplemental note 2). (**F**) The ability of CTCF to form stable, dimeric bridges may also explain how monomeric cohesin halts at two convergent CTCF molecules (each with N-termini encountering cohesin) despite possessing a single identified binding site.

While CTCF-mediated bridges and loops have long been hypothesized,^31,56^ this study provides direct observations of that ability. By adapting a novel configuration of the quad-trap optical tweezers, we developed a methodology capable of separately quantifying both bridge rupture and sliding forces (Figure 5D-E). The CTCF bridges we observed form readily and slide with little resistance yet paradoxically resist rupture forces beyond the stability of dsDNA. Although focus in the literature has shifted to cohesin activity, particularly strong evidence for loop extrusion- independent CTCF bridging exists in the formation of inter-chromosome contacts mediated by CTCF that cannot be explained by expanding loops.^32,33^

The LEM is relatively minimal, incorporating only DNA, cohesin and CTCF, with the mechanisms of CTCF remaining largely uncharacterized.^9^ Our findings, particularly of CTCF’s ability to form bridges independently, provide a unifying explanation for several unresolved phenomena linked to CTCF’s roles in genome organization. First, cohesin loop extrusion is halted by CTCF,^12^ at least in part due to interaction between the two proteins,^27^ with a recent single-molecule study indicating that fragments of the CTCF N-terminus impede extrusion by cohesin.^28^ However, the mechanism by which cohesin is halted at two convergent CTCF molecules remains unclear, as only a single CTCF-interaction site on cohesin has been identified. While the first CTCF encountered may halt cohesin extrusion in one direction, we propose that continued extrusion in the opposite direction reels in a second, convergent CTCF, “zipping” them together and driving bridge formation that would terminate extrusion (Figure 5F). Second, CTCF contributes additional stability to *in vivo* loops, with additional CTCF acting cooperatively^40^ and CTCF depletion reducing loop lifetime.^57^ While it has been speculated that this may be due to CTCF increasing cohesin strength, we propose a simpler explanation - direct bridging by CTCF. Beyond loop extrusion, several recent studies have identified a mitotic role for CTCF at centromeres.^55,58^ One study found that CTCF depletion results in a weakened pericentromeric chromatin spring that is less resistant to tension.^54^ Again, direct CTCF bridging presents a simple explanation, especially with our finding that CTCF bridges resist forces similar to those expected to occur at the centromere during mitosis.

These findings expand our understanding of how CTCF functions in genome organization. Beyond serving as a directional barrier to loop extrusion, CTCF directly modulates DNA mechanics, stabilizes long-range DNA interactions, and forms force-resistant bridges that maintain the ability to be repositioned. The ability to both slide and withstand rupture suggests a flexible, adaptive CTCF:DNA complex capable of accommodating dynamic chromatin rearrangements while maintaining loop integrity. Together, these findings support a model in which CTCF plays an active mechanical role in genome organization rather than solely as a passive barrier to loop extrusion, by forming dynamic, force-resistant DNA bridges that stabilize chromatin loops. These findings provide a foundation for future work integrating additional essential factors, particularly cohesin, to determine how their interplay with CTCF contributes to active chromatin organization and dynamics.

## RESOURCE AVAILABILITY

## Lead contact

Request for further information and resources should be directed to and will be fulfilled by the lead contact, Gijs Wuite

## Data and code availability

- Source and supporting data generated during this study will be available from Zenodo upon publication.
- All original code has been deposited on Zenodo and will be publicly available as of the date of publication.
- Raw optical tweezers data and any additional information required to reanalyze the data reported in this paper are available by request to the lead contact.

## ACKNOWLEDGMENTS

We thank E.J.G. Peterman, M. Spies, D.M. Benson, M. Marzin and members of the Wuite lab for productive discussions and support; and X. Seymonson for eWLC analysis script support. This work was supported by the Netherlands Organization for Scientific Research (NWO) Science (ENW) grant (OCENW.GROOT.2019.012) and Gravitation grant (BaSyC) as well as the European Research Council under the European Union’s Horizon 2020 research and innovation program grant (MONOCHROME, grant agreement no. 883240) to G.J.L.W.

## AUTHOR CONTRIBUTIONS

C.C.C. conceived the study. C.C.C., A.S.B., and G.J.L.W. developed the experimental approach. C.C.C. and N.F.Z. performed protein purification. C.C.C. and M.S. performed optical tweezers experiments. A.R. carried out AFM experiments and analysis. C.C.C. performed data analysis and generated figures. B.J.H. and C.P.B. estimated expected friction. C.C.C. and A.S.B. wrote the manuscript. G.J.L.W. led the research and supported the interpretation of the results. All authors took part in reviewing the content critically and approving the manuscript.

## SUPPLEMENTAL INFORMATION

Document S1. Supplemental notes, Figures S1-S5, and supplemental references

**Video S1. CTCF forms bridges that readily slide along DNA at low applied force, related to Figure 4**

Quad-trap optical tweezers with widefield fluorescence microscopy show CTCF (green) forms a stable bridge at the intersection of two DNA molecules. The CTCF-mediated bridge slides along DNA in the direction of applied force while non-bridging DNA-bound CTCF is readily displaced.

**Video S2. CTCF bridge rupture and separation of bridged dsDNA molecules, related to Figure 4**

Quad-trap optical tweezers with widefield fluorescence microscopy show that CTCF (green) forms a bridge at the intersection of two DNA molecules that ruptures under increasing applied force, leading to separation of the DNA strands. Bridge rupture prior to DNA breaking is rare, with most DNA rupturing while CTCF bridges remain intact (Video S3).

**Video S3. dsDNA breaking while the CTCF bridge remains intact, related to Figure 4**

Quad-trap optical tweezers with widefield fluorescence microscopy show that CTCF (green) forms a bridge at the intersection of two DNA molecules that persists under increasing applied force beyond the stability of DNA. Following DNA breaking, discontinuous DNA segments remain held together by the intact CTCF bridge.

## METHODS

### Expression, purification, and labelling of CTCF

Full-length human CTCF with an N-terminal 6X polyhistidine tag was codon optimized and inserted into the pET28a plasmid (GenScript). Rosetta(DE3)pLysS *Escherichia coli* containing the pET28a-CTCF plasmid were grown in Luria broth at 37°C until reaching OD_600_ = 0.6-0.7 then induced with 1 mM IPTG. 0.5 mM ZnCl_2_ was added to the cells which were then grown for 12 hours at 16°C. Cells were harvested by centrifugation and resuspended in lysis buffer (50 mM HEPES pH 7.5, 750 mM NaCl, 40 mM imidazole, 1 mM DTT, 0.1% Triton X-100, 5% glycerol, 1 mg/ml lysozyme, and protease inhibitor cocktail). Sonication was used to lyse the cells and the lysate was clarified using centrifugation. Clarified lysate was applied to a HisTrap HP column (Cytiva), rinsed with buffer NW (50 mM HEPES pH 7.5, 500 mM NaCl, 40 mM imidazole, and 1 mM DTT), and eluted over a gradient with buffer NE (50 mM HEPES pH 7.5, 500 mM NaCl, 500 mM imidazole, and 1 mM DTT). Fractions containing CTCF were pooled and diluted in buffer H0 (50 mM HEPES pH 7.5, 1 mM DTT) to obtain a NaCl concentration of 150 mM. The diluted fractions were then applied to a HiTrap Heparin HP column (Cytiva), washed with HA buffer (50 mM HEPES pH 7.5, 150 mM NaCl, and 1mM DTT), and eluted over a gradient with HB buffer (50 mM HEPES pH 7.5, 2 M NaCl, and 1mM DTT). Fractions containing CTCF were pooled and concentrated in S buffer (50 mM HEPES pH 7.5, 300 mM NaCl, and 1mM DTT). Concentrated fractions were applied to a HiPrep 26/60 S-400 column (Cytiva) and eluted. Fractions containing purified CTCF were exchanged into CTCF storage buffer (50 mM HEPES pH 7.5, 300 mM NaCl, 1mM DTT, and 20% glycerol) and stored at -70°C. Purity was verified by SDS-PAGE analysis.

For use in confocal and widefield imaging, CTCF was fluorescently labelled with Alexa Fluor 555. CTCF was dialyzed in labelling buffer (50 mM HEPES pH 7.0, 300 mM NaCl, 1mM DTT, and 20% glycerol). Alexa Fluor 555 NHS Ester (Invitrogen) was added to the protein to yield an 8-fold molar excess. The reaction was incubated at 4°C while rotating in darkness for 12 hours. Excess dye was removed and buffer exchanged to CTCF storage buffer using a Zeba Spin Desalting Column (ThermoScientific). Labelling efficiency was calculated using the absorbance at 280 nm and 555 nm to determine the ratio of Alexa Fluor 555 to CTCF molar concentrations. Aliquots were flash frozen and stored at -70°C.

### Dual-trap optical tweezers

Experiments were performed using a commercial dual-trap optical tweezers setup (C-Trap, LUMICKS) equipped with confocal fluorescence microscopy. Two optical traps were generated using a 1064 nm laser (4 watts). A u-Flux microfluidic system (LUMICKS) was connected to a five-channel flow cell. The preparation of end-biotinylated bacteriophage λ-DNA was carried out as described previously^59^. Streptavidin-coated polystyrene beads (3.45 μm diameter, Gentaur) were trapped in the first channel, and λ-DNA with biotin-labelled ends was tethered between two beads by capturing the DNA in the next channel. Tethered DNA was transferred to a channel containing CTCF Reaction Buffer (20 mM HEPES pH 7.5, 25 mM NaCl, 1 mM MgCl_2_, 1 mM DTT, 0.1% TWEEN) and single tether formation was verified by stretching the DNA to overstretching forces (∼65 pN) and confirming characteristic force-extension behavior. All experiments were carried out in CTCF Reaction Buffer, unless otherwise stated.

### Confocal fluorescence imaging quantification of CTCF binding

Tethered λ-DNA molecules were transferred to a channel containing CTCF^555^ in concentrations ranging from 500 pM to 50 nM CTCF and allowed to incubate for one minute. These experiments were conducted under 20 pN of DNA tension; while this does not have an effect on the coating, this is a sufficient force to prevent loop formation, which could result in DNA bridging. Confocal fluorescence excitation was generated using a 532 nm laser at 10.8 mW. Images of CTCF^555^ associated with tethered DNA were captured using confocal scanning. ImageJ was used to quantify the total fluorescence signal associated with DNA and subtract background fluorescence. For each concentration, total fluorescence values from at least five tethers (*N* = 10, 10, 9, 10, 9, 7, 5, and 9) were averaged and plotted against CTCF concentration. To determine the binding affinity and cooperativity of CTCF interaction with long, non-specific dsDNA, fluorescence data were analyzed using the cooperative nearest-neighbor model of McGhee and von Hippel.^37^ The fractional binding density per base pair (ν) is related to the concentration of free CTCF (*L*) using the following equation

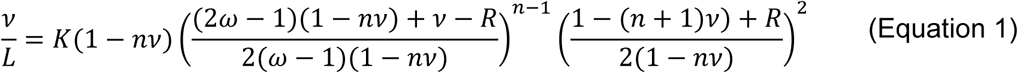

Where

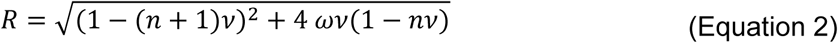

Here, *K* is the equilibrium binding constant (nM^-1^), *n* is the binding site footprint in base pairs, and ω is the nearest-neighbor cooperativity factor. The binding footprint of 29 ± 1.6 bp was determined using the measured intensity of a single monomer (Figure S2) to calculate the total number of bound monomers on the length of tethered dsDNA. Total fluorescence was normalized to determine fractional coverage, θ, which was related to the parameters above using θ = *n* ν. Parameters *K* and ω were determined by nonlinear least squares minimization using MATLAB.

### Measuring impact of CTCF binding on DNA force response

To evaluate how CTCF alters the mechanical properties of dsDNA, λ-DNA was tethered between two optically trapped beads in the C-Trap setup. After confirming proper tethering and canonical dsDNA force-extension behavior, three consecutive force-extension curves were recorded for bare dsDNA at a pulling speed of 0.5 μm/s. The tethered dsDNA was then transferred into a channel containing 50 nM CTCF and incubated for one minute to allow saturation. Subsequent force-extension curves were acquired for the CTCF-coated DNA under identical conditions. Bare dsDNA and CTCF-coated dsDNA curves were fit to the extensible worm-like chain (eWLC) model to extract persistence length (L_p_), contour length (L_c_), and stretch modulus (S) ^60^.

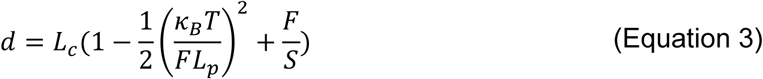

Fits were performed using a custom Python script, and statistical significance of mechanical parameters was determined using unpaired t-tests performed with GraphPad Prism.

### Observation of CTCF diffusion

To observe diffusion of CTCF on non-specific DNA, λ-DNA was tethered between beads in the C-Trap and transferred to a channel containing ≤1 nM CTCF^555^, conditions under which individual binding and sliding events were observed. Confocal fluorescence excitation was generated using a 532 nm laser at 10.8 mW. Kymographs were generated by collecting consecutive confocal line scans along the axis of the DNA using a pixel size of 75 nm and pixel dwell time of 1.5 ms. A previously employed custom MATLAB program^61^ was used to analyze kymographs and yield time, position, and fluorescence intensity information for individual particle trajectories. Monomeric CTCF trajectories were selected based upon single-step photobleaching and intensity (Figure S2). Monomers with unconstrained sliding trajectories were selected for further diffusion analysis. MSD was calculated from kymograph trajectories, using time (t) and position (x):

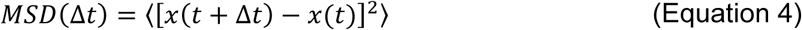

MSD was plotted as a function of time lag and the diffusion constant was calculated from the slope using a custom MATLAB script. In addition to unconstrained mobile monomers, the sliding trajectories of CTCF monomers that exhibited more complex diffusive behavior, including pausing, and CTCF oligomers were similarly analyzed to yield individual diffusion constants.

### Quad-trap optical tweezers

Quad-trapping experiments were performed using a commercial quad-trap optical tweezers setup (C-Trap, LUMICKS) equipped with widefield fluorescence microscopy. Four independently steerable optical traps were generated using a 1064 nm laser. The microfluidic system, channel arrangement, and DNA constructs are consistent with those used in dual-trapping experiments. Slightly smaller streptavidin-coated polystyrene beads (3.13 μm diameter, Gentaur) were used in quad-trapping experiments.

Following capture and tethering of two parallel dsDNA molecules, the second strand was lowered in the Z plane and rotated to be perpendicular to the first strand. The second strand was then translocated underneath the first strand, creating a “+” configuration. The second strand was then brought into alignment in the Z plane with the first strand, generating a single point of contact at the intersection of the two dsDNA molecules. ∼10 pN of tension was applied to each molecule. The crossed dsDNA was transferred to a channel containing 25 nM CTCF^555^ and allowed to incubate for one minute. Following incubation, the crossed DNA molecules were transferred back to a buffer-only channel and tested for bridging. Illustrations of bead orientation and manipulations to generate and probe bridges are shown in Figures S4 and S5.

### Observation of CTCF sliding bridges

Following formation of a CTCF-mediated bridge between the crossed dsDNA molecules, the bridges were probed with force to generate sliding. While the horizontal strand was left stationary, the vertical strand was translocated along the horizontal strand. Bridge sliding could also be generated by moving the vertical strand along its own axis, though interpretation of the forces generated in this orientation is less reliable due to movement of the second bead on which force is monitored. Detailed illustration of dsDNA manipulation to form and probe bridges is found in Figures S4 and S5.

CTCF-mediated bridges observed sliding away from bead 1 were selected. Force on the bridge was determined by subtracting the tension on the horizontal dsDNA molecule from the total force on bead one. Widefield fluorescence microscopy images of the bridged dsDNA were correlated with the measured force. Bead positions and measured forces were captured using LUMICKS Bluelake software while bridge position was analyzed using the ImageJ Manual Tracking plugin.

### Observation of CTCF bridge ruptures

CTCF-mediated bridges were formed using the same procedure as applied to generate sliding bridges. Following bridge formation, the vertical DNA strand was rotated and moved beneath the horizontal strand, resulting in two parallel strands held together only by CTCF bridging. The beads tethering the bottom strand were moved downward to increase separation distance between the top and bottom bead pairs and to increase the force applied to the bridge. The forces at which ruptures occurred were recorded and displayed as a cumulative distribution.

### Atomic force microscopy

pUC-LUMICKS plasmid was digested to yield 2.1kbp linear DNA and purified using an Amicon Ultra Centrifugal Filter. DNA samples were prepared starting from a 2.5 ng/ul aliquot. 36 nM CTCF was added to DNA and incubated. Samples were diluted 5x in mQ water and then a 1ul droplet of sample was spotted together with a 1 ul droplet of NiCl2 to promote adsorption onto freshly cleaved mica. After 5 minutes of equilibration at room temperature, the sample was rinsed with mQ water and then dried with a gentle Ar flow. After this, the sample was measured with the AFM. For the control CTCF sample, a 36 nM aliquot was diluted 10x in mQ water and spotted on freshly cleaved mica. The sample was allowed to dry at room temperature and was then imaged with the AFM. Imaging the CTCF sample in this way, without the use of NiCl2 and rinsing step allows for a clear picture of the pure protein sample, as there are no events potentially influencing the interaction of the protein with the mica surface.

Experiments were carried out using a Bruker Multimode 8, equipped with SNL-10 probes (cantilever C, nominal spring constant 0.24 N/m). Images were performed in air, in PeakForce Tapping mode, optimizing the imaging parameters to accurately follow the sample morphology without introducing any artifact. Results from Figure 3E-F refers to *N* = 16 2×2 mm images for the DNA samples and *N* = 20 2×2 mm images for the DNA-CTCF samples. Images were processed using Gwyddion 2.58. Strand counts were obtained by manually selecting individual strands, while end-to-end distance distributions were obtained from Gwyddion 2.58 after manually pinpointing the ends of each strand. Only isolated strands, whose topology could be clearly traced from one end to the other, were taken into consideration for the quantitative analyses (strand count and end-to-end distance distributions).

### Quantification and statistical analysis

Statistical analysis was performed using GraphPad Prism (v.10.1.4) or specified custom MATLAB (v2024b) and Python (v.3.13.2) scripts. To compare properties of DNA with and without bound CTCF (Figure 2I-L), unpaired t-tests were performed. Exact P values and replicate numbers are described in the corresponding figure legend. Unless otherwise indicated, all data is reported as mean ± S.E.M. No statistical methods were used to determine sample sizes. All experiments were replicated successfully on multiple days using freshly prepared samples and buffers to ensure reproducibility of results. Plots shown in figures were generated using GraphPad Prism.

