## Supplemental Information for "Dynamic CTCF bridges drive genome organization"

### Supplemental Note 1: Estimating the characteristic energy barrier for the CTCF-dsDNA interaction

The observed 1D diffusion constant $D_{\mathrm{obs}}$ of a protein sliding along a DNA molecule can be expressed as

$$D_{\mathrm{obs}} = D_{\mathrm{hyd}} * e^{-\left(\frac{\sigma}{kT}\right)^2}. \qquad \text{(Equation 5)}$$

Here, $D_{\mathrm{hyd}}$ is the hydrodynamic diffusion constant and the exponential factor accounts for the rough energy landscape of the protein-DNA interactions along the DNA, where $\sigma$ denotes the typical energy barrier height that the protein must overcome when hopping between binding sites [S1]. To assess the energy potential of CTCF interactions with non-specific DNA, we thus need to know the observed diffusion constant ($D_{\mathrm{obs}}$), as well the expected diffusion constant without CTCF-dsDNA binding interactions ($D_{\mathrm{hyd}}$).

From the Einstein-Smoluchowski relation, we can relate the hydrodynamic diffusion constant of a particle to the thermal energy $kT$ and the Stokes drag $\zeta$ through

$$D_{\mathrm{hyd}} = \frac{kT}{\zeta}. \qquad \text{(Equation 6)}$$

For 1D translational diffusion along the DNA strand, the drag scales linearly with particle size. For a spherical particle of radius $R$, it is given by $\zeta_{\mathrm{tra}}^{\mathrm{sp}} = 6\pi\eta R$, with $\eta \approx 10^{-3}\mathrm{kg\,m^{-1}s^{-1}}$ being the viscosity of water. In addition to translational diffusion, proteins who search for a specific target sequence follow the DNA helical pitch, therefore rotating and exhibiting an additional rotational drag. This often results in much slower diffusion, since the rotational drag $\zeta_{\mathrm{rot}}^{\mathrm{sp}}$ essentially scales with the cube of the size [S2]; e.g., again for a spherical particle

$$\zeta_{\mathrm{rot}}^{\mathrm{sp}} = (8\pi\eta R^3 + 6\pi\eta R(R+r)^2)\left(\frac{2\pi}{l}\right)^2. \qquad \text{(Equation 7)}$$

Here, the first term corresponds to the intrinsic rotational drag of a sphere around its own axis, and the second term arises from the particle rotating at an external axis. In our case, this external axis is equal to the radius of dsDNA ($r = 1\mathrm{nm}$) together with the particle radius [S3]. The final term is a geometric conversion factor which translates the proteins linear motion to the corresponding angular rotation around the helix, with $l \approx 3.4\mathrm{nm}$ constituting the helical pitch of the DNA. Because the rotational drag scales cubically with size, for all our considered protein sizes we find that the rotational drag dominates ($\zeta_{\mathrm{rot}} \gg \zeta_{\mathrm{tra}}$) and thus ignore the translational drag.

Diffusing proteins are often approximated as spherical particles [S4,5], but this simplification may be inadequate for CTCF. The known crystal structures of CTCF [S6] show elongated zinc finger domains along the DNA sequence; we therefore assume that a cylinder of length $L_{\mathrm{zf}} \approx 11\mathrm{nm}$ and radius $R_{\mathrm{zf}} \approx 1.5\mathrm{nm}$ - both estimated from existing crystal structures - gives a more faithful approximation of the protein core. In addition, CTCF features a sizeable disordered region on both N- and C-termini, which we approximate as spheres. For the radii, we assume the radius of gyration $R_g$ which for unstructured proteins of sequence length $N$ has been estimated by $R_g = \sqrt{Nbl/6}$, with assumed value of $b \approx 0.38\mathrm{nm}$ and $l \approx 0.8\mathrm{nm}$ [S7]. Plugging in the sequence lengths of $N_{\mathrm{N}} = 265$ and $N_{\mathrm{N}} = 150$ for the N- and C-terminus, we find radii of $R_{\mathrm{N}} \approx 3.7\mathrm{nm}$ and $R_{\mathrm{C}} \approx 2.8\mathrm{nm}$.

To calculate the total hydrodynamic drag, we need to determine the contribution from both the protein core and the disordered regions at the termini. We approximate the protein core as a cylinder [S8] of length $L$ and radius $R$ and determine the rotational drag with

$$\zeta_{\mathrm{rot}}^{\mathrm{cy}} = 4\pi\eta R^2 L\left(\frac{2\pi}{l}\right)^2. \quad \text{(Equation 8)}$$

The two termini are modelled by spheres of $R_{\mathrm{N}} = 2.8$ and $R_{\mathrm{C}} = 3.7\mathrm{nm}$, located at an offset of $r = 1\mathrm{nm}$ from the rotational axis, identical to the model of Bagchi *et al* [S3]. Similar to the spherical case, the translational drag of the cylindrical section contributes little to the overall drag, so the total hydrodynamic drag becomes

$$\zeta_{\mathrm{tot}}^{\mathrm{CTCF}} \approx \zeta_{\mathrm{N}}^{\mathrm{CTCF}} + \zeta_{\mathrm{C}}^{\mathrm{CTCF}} + \zeta_{\mathrm{cil}}^{\mathrm{CTCF}}$$
$$\approx \left(\frac{2\pi}{l}\right)^2 \left(\left(8\pi\eta {R_{\mathrm{N}}}^3 + 6\pi\eta R_{\mathrm{N}}(R_{\mathrm{N}} + r)^2\right) + \left(8\pi\eta {R_{\mathrm{C}}}^3 + 6\pi\eta R_{\mathrm{C}}(R_{\mathrm{C}} + r)^2\right) + 4\pi\eta R_{zf}^2 L_{zf}\right) \quad \text{(Equation 9)}$$

We therefore arrive at an approximate hydrodynamic drag for CTCF $\zeta_{\mathrm{tot}}^{\mathrm{CTCF}} \approx 1.52 * 10^{-8}\mathrm{kg/s}$, and through Eq. 5 at a hydrodynamic diffusion constant of $D_{\mathrm{hyd}}^{\mathrm{CTCF}} \approx 2.7 * 10^{-13}\mathrm{m}^2/\mathrm{s} = 0.27\mathrm{\mu m}^2/s$. This value is much smaller than the hypothetical hydrodynamic diffusion constant of a similar shaped protein which does not rotate, which is

$$D_{\mathrm{hyd}}^{\mathrm{tra}} = \frac{kT}{\zeta_{\mathrm{tra}}^{\mathrm{p}}} = \frac{kT}{\zeta_{tra}^{sp,N} + \zeta_{tra}^{sp,C} + \zeta_{tra}^{cyl}} \approx \frac{26\ \mathrm{\mu m}^2}{\mathrm{s}}. \quad \text{(Equation 10)}$$

Here, we used the following [S9], $\zeta_{\mathrm{tra}}^{\mathrm{sp}} = 6\pi\eta R$, and $\zeta_{tra}^{cyl} = 2\pi\eta L_{zf}/(\ln\frac{L_{zf}}{R_{zf}} + \gamma)$ with $\gamma = -0.08$.

Next, we can use our derived hydrodynamic diffusion constant for rotating CTCF to determine the characteristic energy barrier associated with the non-specific diffusion interaction of CTCF, which yields

$$\sigma_{\mathrm{CTCF}} = \sqrt{\ln\left(\frac{D_{\mathrm{hyd}}^{\mathrm{CTCF}}}{D_{\mathrm{obs}}^{\mathrm{CTCF}}}\right)}\, kT \approx 1.04kT. \quad \text{(Equation 11)}$$

Note that this is very similar to energy barriers reported for many other proteins showing facilitated diffusion [S4,5].

#### Supplemental Note 2: Deducing whether the observed dragging force is compatible with the diffusive motion of the protein

In the previous section, we derived a characteristic energy barrier of $\sigma_{\mathrm{CTCF}} \approx 1.04kT$ from the observed 1D diffusion constant. Using this, we can determine the expected dragging force when a bridged DNA strand is pulled past another strand. Compared to the case above, where we considered a static piece of dsDNA on which a mobile protein diffuses, we now face the opposite configuration. The bridged protein that slides along the strand is fixed to the other strand, and is therefore unable to rotate. Thus, if the protein still follows the helical pitch during the dragging, then the dsDNA strand must rotate instead, which is possible since the dsDNA is not torsionally constrained. As a result, we need to include a hydrodynamic drag arising from the rotating dsDNA. Using Eq. 8 and modelling the DNA as a cylinder of radius $r$ and length $L_{\mathrm{DNA}}$, we can write the drag as

$$\zeta_{\mathrm{hyd}}^{\mathrm{DNA}} = 4\pi\eta r^2 L_{\mathrm{DNA}}\left(\frac{2\pi}{l}\right)^2. \quad \text{(Equation 12)}$$

The energy landscape from the CTCF-dsDNA interaction remains the same, so analogous to Eq. 5 we expect an effective drag of

$$\zeta_{\mathrm{eff}}^{\mathrm{DNA}} = \zeta_{\mathrm{hyd}}^{\mathrm{DNA}} * e^{\left(\frac{\sigma}{kT}\right)^2}. \qquad \text{(Equation 13)}$$

Here, we ignore the translational drag of the bridge as we assume it to be negligible. Note that the exponential factor appears with opposite sign compared to Eq. 5, due to the diffusion and friction being inversely related (Eq. 6). This allows us to determine the expected drag force using the Stokes relation $F_{\mathrm{drag}}^{\mathrm{DNA}} = \zeta_{\mathrm{eff}}^{\mathrm{DNA}} * v_{\mathrm{drag}}$, where $v_{\mathrm{drag}} \approx 0.5\ \mu m/s$ is the observed speed of the bridge while dragging. The length of the rotating dsDNA is at most equivalent to the total length of tether dsDNA (16.5 $\mu m$), though nicks could result in shorter rotating segments. For the upper limit of the dragging force, we then find

$$F_{\mathrm{drag}}^{\mathrm{DNA}} = 4\pi\eta r^2 L_{\mathrm{DNA}} \left(\frac{2\pi}{l}\right)^2 e^{\left(\frac{\sigma}{kT}\right)^2} v_{\mathrm{drag}} = 16\pi^3\eta \frac{r^2}{l^2} L_{\mathrm{DNA}} * e^{\left(\frac{\sigma}{kT}\right)^2} v_{\mathrm{drag}} \approx 1.05\ \mathrm{pN}, \qquad \text{(Equation 14)}$$

which is in agreement with the upper limit for the measured dragging force.

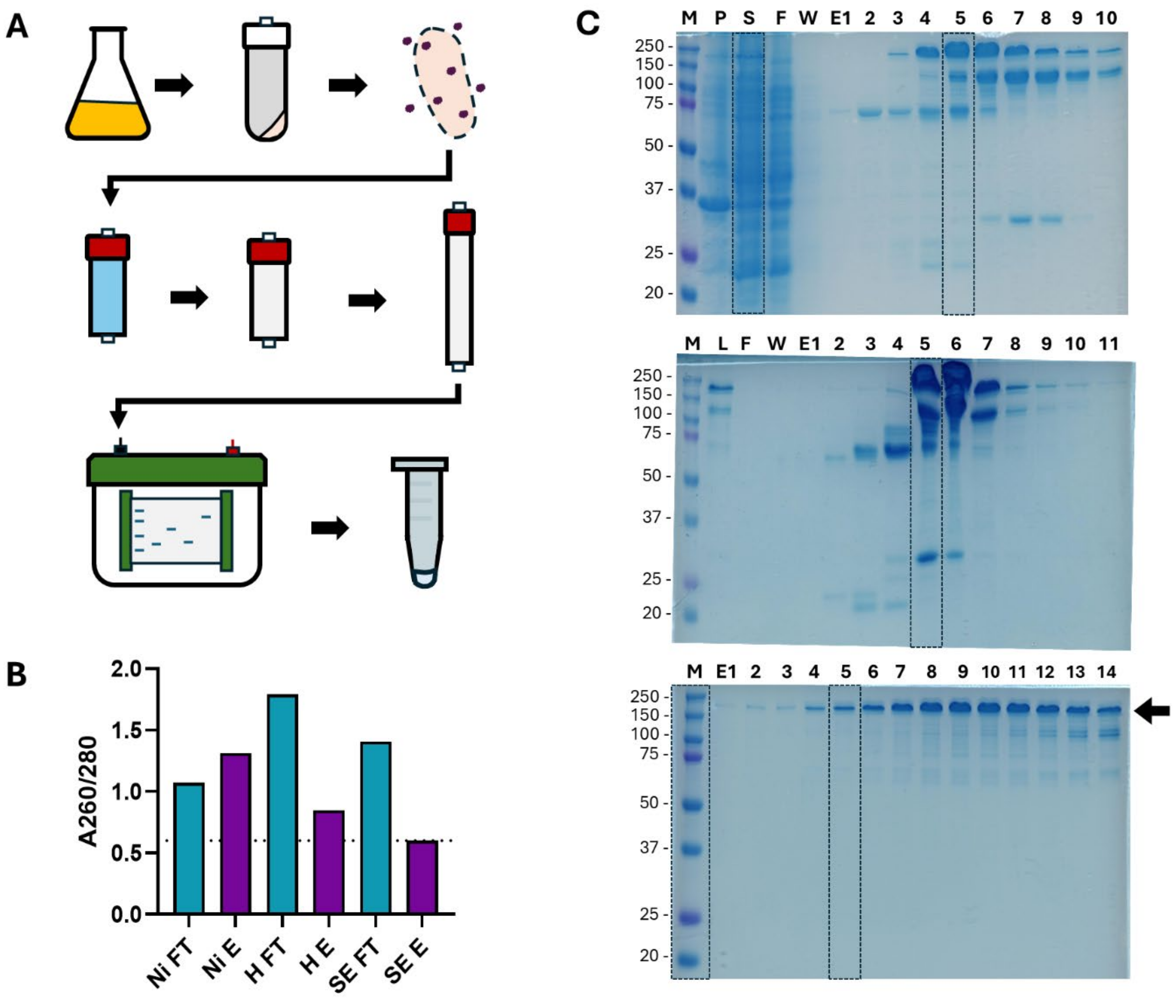


**Figure S1. Purification of full-length CTCF. Related to Figure 1.** (**A**) Schematic overview of protein purification workflow. Full-length human CTCF with an N-terminal 6×His tag was expressed in *E. coli* Rosetta(DE3)pLysS and grown in Luria broth with 0.5 mM $ZnCl_2$. Expression was induced with 1 mM IPTG at $OD_{600}$ = 0.6–0.7 and carried out overnight at 16 °C. Cells were harvested, lysed by sonication, and clarified by centrifugation. CTCF was purified using a HisPrep nickel ($Ni^{2+}$) affinity column, HiTrap Heparin column, and HiPrep 26/60 S-400 size-exclusion column. Protein purity was characterized via SDS-PAGE. The final protein product was stored at −70 °C in storage buffer (50 mM HEPES pH 7.5, 300 mM NaCl, 1 mM DTT, 20% glycerol). (**B**) $A_{260}/A_{280}$ ratios for elution (E) and flowthrough (FT) fractions from each chromatography step, nickel (Ni), heparin (H), and size exclusion (SE), were measured. Elimination of nucleic acid contamination was only reached following the final size exclusion step. (**C**) SDS-PAGE analysis of purification: (top) nickel affinity, (middle) heparin affinity, and (bottom) size exclusion. Lane markers indicate (M) molecular weight ladder, (P) insoluble pellet, (S) soluble supernatant, (F) flow through, (W) wash, and (E#) elution fractions. Arrow indicates CTCF on the final gel. Boxes indicate data shown in Fig. 1D.

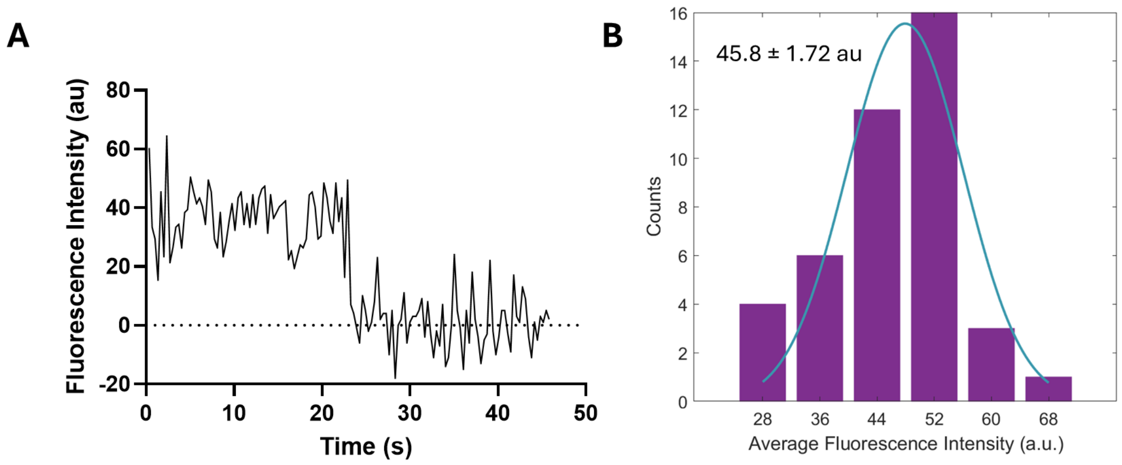


**Figure S2. Determination of fluorescence intensity of CTCF monomers. Related to Figure 2.** (**A**) Single-step photobleaching events were analyzed to determine the intensity of a single CTCF monomer labelled with AlexaFluor555. (**B**) Collected photobleaching steps (*N* = 42) show a regular distribution with an average monomer intensity of 45.8 ± 1.72 au.

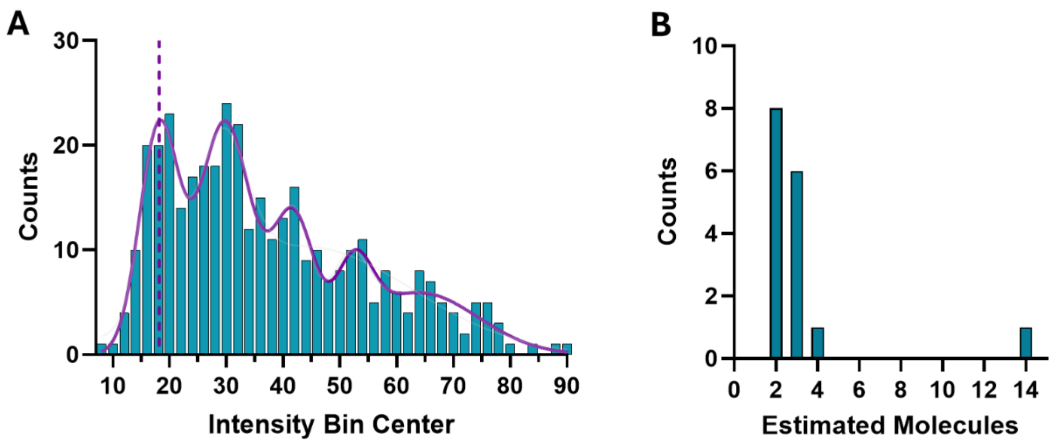


**Figure S3. Quantification of CTCF molecules associated with bridges. Related to Figure 4.** (**A**) Intensity peaks associated with CTCF bound to DNA in bridging experiments were extracted from fluorescence images, corrected for background fluorescence and fit to a sum of five Gaussian distributions (purple line). The single fluorophore intensity was determined to be 18.1 ± 3.4 au. (**B**) Bridge intensities were calculated by averaging values across three frames. The estimated number of molecules was determined by dividing the average intensity by the single fluorophore intensity, with the majority of bridges estimated to be composed of CTCF dimers. Notably, no bridges were found with only a single CTCF molecule.

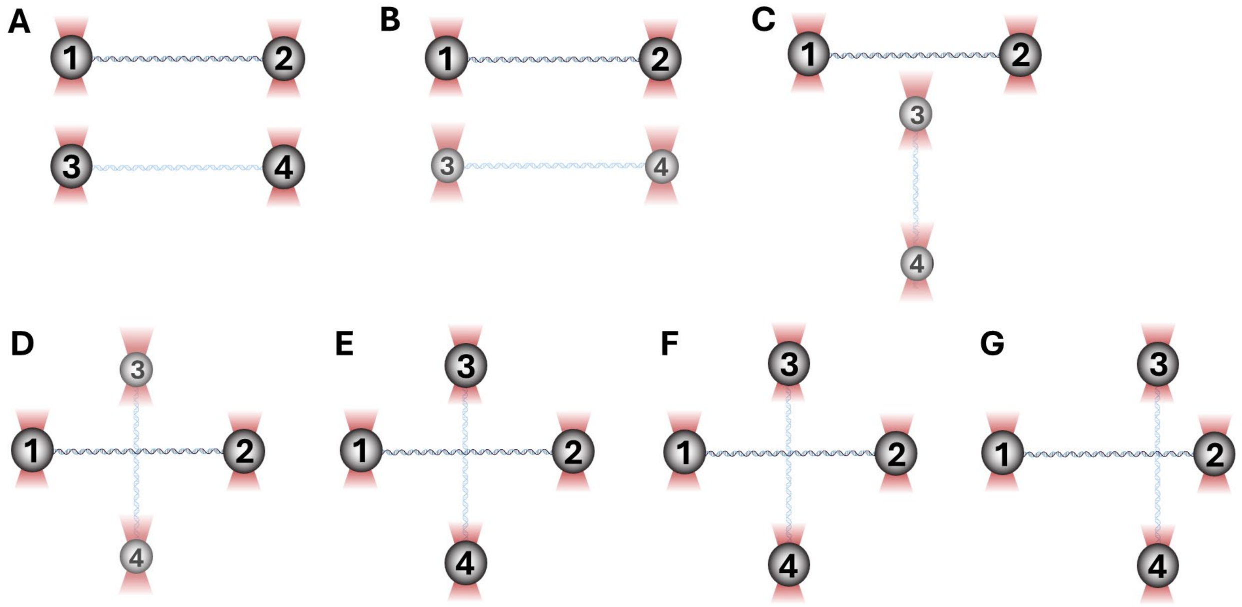


**Figure S4. Illustration of CTCF-mediated bridge formation and force induced sliding. Related to Figure 4.** (**A**) Four beads are optically trapped and DNA is tethered between beads 1 and 2 and beads 3 and 4. A single tether is confirmed by verifying typical force-extension behavior. (**B**) Beads 3 and 4 are lowered in the Z plane to allow clearance below beads 1 and 2. (**C**) Beads 3 and 4 are rotated so the tethered strand is perpendicular to the strand between beads 1 and 2. (**D**) Bead 3 and 4 are moved simultaneously so bead 3 moves below the molecule tethered between bead 1 and 2. (**E**) Beads 3 and 4 are returned to the same Z position as bead 1 and 2, bringing the two tethered stands into contact at a single point. Equivalent tension is applied to both strands (5-10 pN). (**F**) The crossed molecules are incubated in a channel containing 20 nM $CTCF^{555}$ for 1 minute. The crossed DNA molecules are then returned to a channel containing only CTCF reaction buffer. (**G**) To probe for bridging and force induced sliding, Beads 3 and 4 are moved simultaneously along the axis of the molecule tethered between beads 1 and 2.

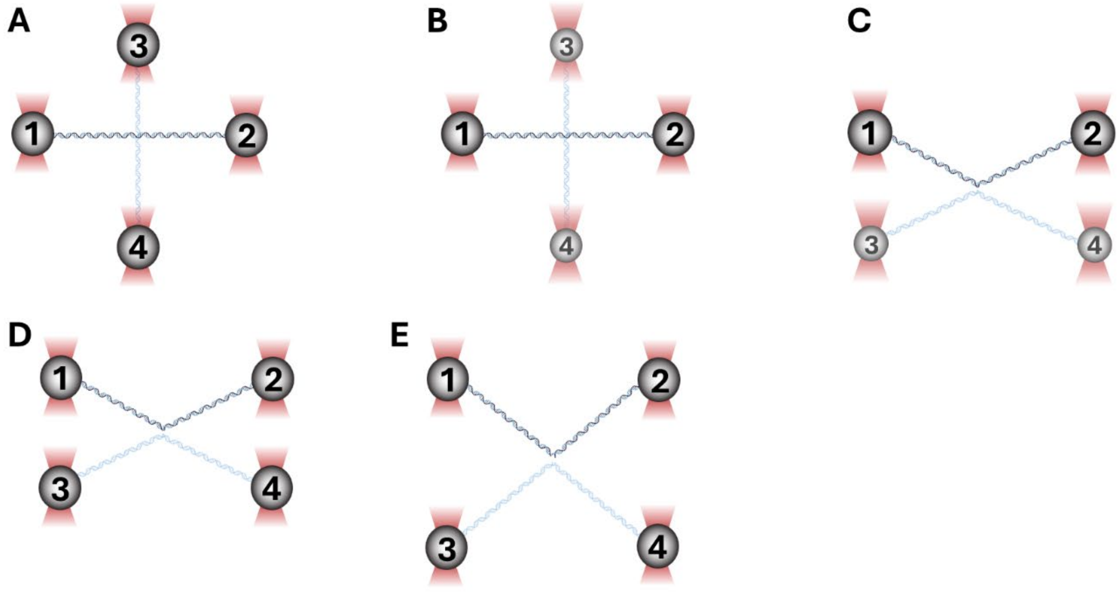


**Figure S5. Illustration of CTCF-mediated bridge force probing for ruptures.** (**A**) The CTCF-mediated bridge is formed as in Figure S4. (**B**) Beads 3 and 4 are lowered in the Z plane to allow clearance under the molecule tethered between beads 1 and 2. (**C**) Bead 3 is moved below the horizontal strand. Beads 3 and 4 are relocated to orient the DNA molecules in a parallel orientation. (**D**) Beads 3 and 4 are returned to the same Z plane as beads 1 and 2. (**E**) Beads 3 and 4 are simultaneously moved downward to apply force to the CTCF-mediated bridge.
